# Navigational systems in the human brain dynamically code for past, present, and future trajectories

**DOI:** 10.1101/2023.08.22.554387

**Authors:** You (Lily) Cheng, Sam Ling, Chantal E. Stern, Elizabeth R. Chrastil

## Abstract

Navigational trajectory planning requires the interaction of systems that include spatial orientation and memory. Here, we used a complex navigation task paired with fMRI pattern classification to examine head and travel direction tuning throughout the human brain. Rather than a single, static network, we report multiple simultaneous subnetworks that 1) have strong connections with both allocentric (world-centered) and egocentric (viewer-centered) movement trajectories, 2) change during the course of exploration, 3) code for past and future movements as well as the present direction, and 4) are strongest for individuals who convert their trajectories into egocentric movements once they have learned the environment. These findings shift our understanding of the neural processes underlying navigation from static structure-function relationships to a dynamic understanding of the multiple brain networks that support active navigation. The insights into the nature of individual navigation abilities uncovered here challenge the dominant framework of largely allocentric coding for successful navigation in complex environments, and replace this with a new framework that relies on multiple co-existing dynamic computations.

## Introduction

Head and travel directions are crucial in human wayfinding. The fundamental signals of head direction cells have been found in rodents^1–3^, including evidence for both allocentric (world-centered) and egocentric (viewer-centered) coordinate frames during movement. For example, hippocampal place cells and theta oscillations reflect past and future path trajectories^4–8^. Moreover, retrosplenial cortex and posterior parietal cortex code for routes and egocentric relationships^9–11^.

In this study, we examined whether travel direction signals can be decoded in the human brain during active navigation, and if so, how individual decoding discriminability relates to navigation performance. We tested the hypothesis that humans possess multiple co-existing reference frames that could be represented at the same time, rather than a single, static allocentric coding. That is, rather than individual brain structures performing individual aspects of navigation (e.g., the hippocampus computes metric cognitive maps), we propose a new, richer framework positing that multiple computations are dynamically performed simultaneously across the human brain. To examine this question, we used multi-voxel pattern classifiers to decode both allocentric (north, south, east, west) and egocentric (left, right, straight) coordinate systems at the present time, as well as past and future trajectory information.

Previous human neuroimaging studies have found brain areas that are sensitive to head direction, including retrosplenial cortex, thalamus, precuneus, extrastriate cortex, and early visual cortex^12–18^. These studies used neural adaptation methods, which do not discriminate one direction from another - yet this discrimination is the hallmark of head direction cells in animals^1,2^. Other studies that have used pattern classification methods relied on open field tasks or static images^13,17,18^. Open field tasks can discriminate between directions (30 vs 60 degree), but cannot be used for specific alignment or for trajectory planning, and static images provide just a single snapshot without the richness of the trajectory of movement. In this study, we sought to examine navigational signals in the human brain while individuals navigated around a complex, dynamic, and naturalistic environment that is more like environments people experience in daily life. In doing so, we sought to investigate the relationship between allocentric and egocentric reference frames in the human brain during navigation trajectory planning.

In an fMRI study, we tested a large group of people (N = 98), who actively navigated in a complex virtual maze environment (Figure 1a, 1b)^19^. The navigation task consisted of an exploration phase and a test phase. During the 16-minute exploration phase, participants freely explored the maze to locate 9 objects, and were instructed to remember their locations. The maze was aligned along with four cardinal directions (arbitrarily defined as north, south, east, and west, see Methods). For each of the 48 test trials, participants started at one object and were instructed to navigate towards another object, within a 45 second time limit (Figure 1c). To reduce feedback during the test phase, all objects were replaced with red spheres, so participants had to go to the location where they remembered the object was. Behavioral performance was measured by the proportion of correct trials in the task; performance ranged from near 0 to 100% accuracy (Figure 1d). This range allowed us to assess the relationship between the accuracy of the classifiers in discriminating directions from brain signals (i.e., classifier strength) and task performance. We also tested the relationship with path efficiency (see Methods, Supplement).

**Figure 1.**
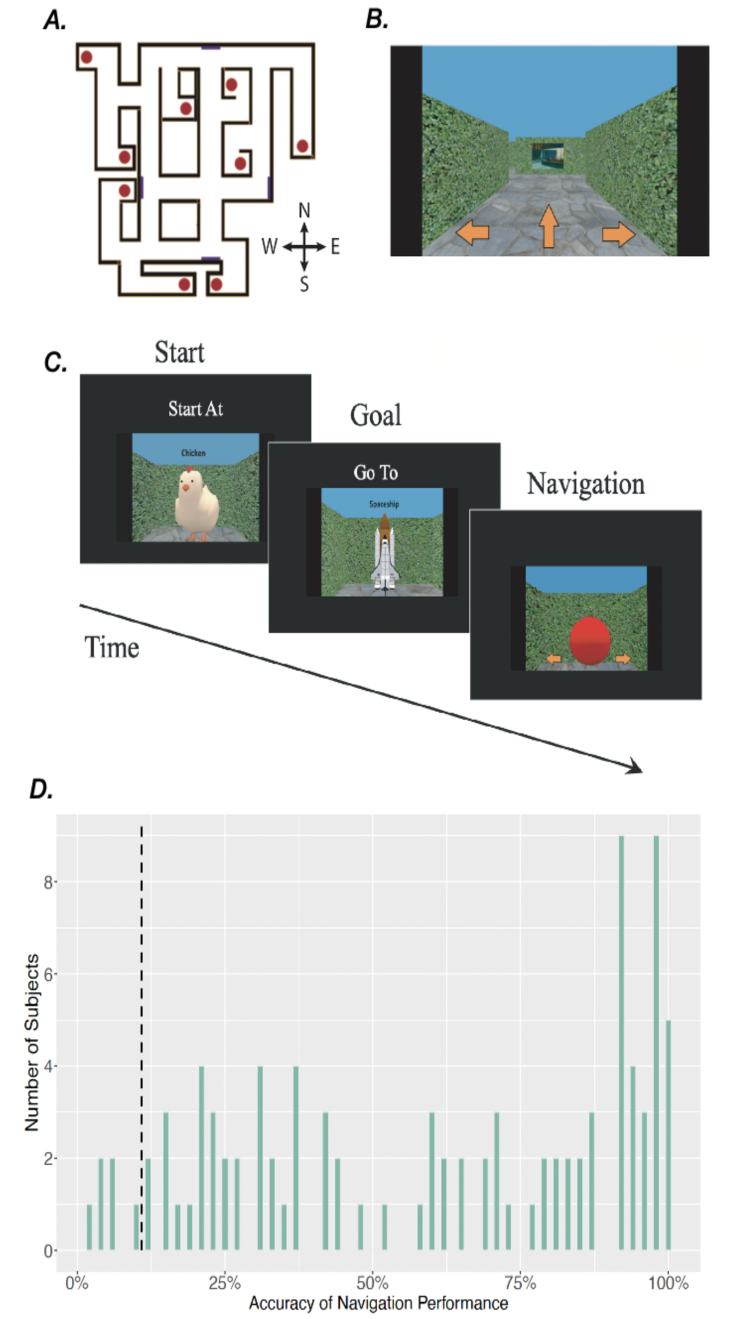
Navigation task and behavior. A) Overhead view of the maze with 9 target objects (red dots) and 4 landmark paintings (purple rectangles) and arbitrary compass directions. Participants never saw this view. B) View from the maze at an intersection. Arrows indicate possible choices (left, right, or straight). C) An example trial from the test phase, which started at one object, going to another object. All objects turned into red spheres to minimize feedback. D) Histogram of accuracy for n=98 participants. Bin size was set to 1%; black dashed line indicates chance performance.

Critically, during both exploration and test phases, participants made choices about where to go next while stationary; they viewed a static image that displayed the possible choices (left, straight, and right) at each intersection (Figure 1b). Movement was gated such that each button press at each choice point caused visual movement to the next choice point (see Methods for a link to video of the task). This static time point and abundant trajectory information allowed us to distinguish between five different classification schemes at any given moment at an intersection: the current allocentric head direction (while stationary), the upcoming allocentric and egocentric movements (next timestep), and the past allocentric and egocentric movements (previous timestep). We were also able to test for allocentric and egocentric classification *during* the movements, to determine how these strong movement cues relate to travel direction signals in the brain.

We conducted intra-subject multivariate pattern classification for different head and travel directions in distributed regions of interest (ROIs) including the hippocampal and parahippocampal cortices^20,21^, basal ganglia^22–24^ (caudate, putamen, nucleus accumbens), visual cortex (extrastriate cortex^16^ and early visual cortex^13,17,25^), thalamus^12^, and parietal lobe (retrosplenial cortex^12,13,17,26^, precuneus^12,15^). These regions have previously been implicated in head direction or egocentric coding, as well as for their general importance to navigation and decision making (see Methods).

## Results

### Allocentric Head and Movement Direction Classifiers

While first exploring and learning the environment, when participants were stationary at an intersection (at the “present” time), we found a network of regions within which we could decode allocentric head direction (Figure 2a, 2e, 2i). Specifically, we found that allocentric head direction information could be decoded in the hippocampus, retrosplenial cortex, thalamus, the striatal regions of caudate and putamen, and early visual cortex, suggesting that head direction signals are also providing information for navigation. Classification performance within this network was highly correlated (r [0.62, 0.89]) between most brain regions (Figure 2e), on any given trial, with an interconnected network relationship (Figure 2i). Yet during the test phase, we could only classify allocentric head direction when stationary in the early visual cortex (Figure 2b). Although correlations between classifications in all ROIs were significant, (r [0.42, 0.92]; Figure 2f), the network analysis revealed that the early visual cortex was largely separate from other regions, with a sub-network of retrosplenial, extrastriate, and nucleus accumbens bridging to the remaining ROIs, which tended to have paired couplings (Figure 2j).

**Figure 2.**
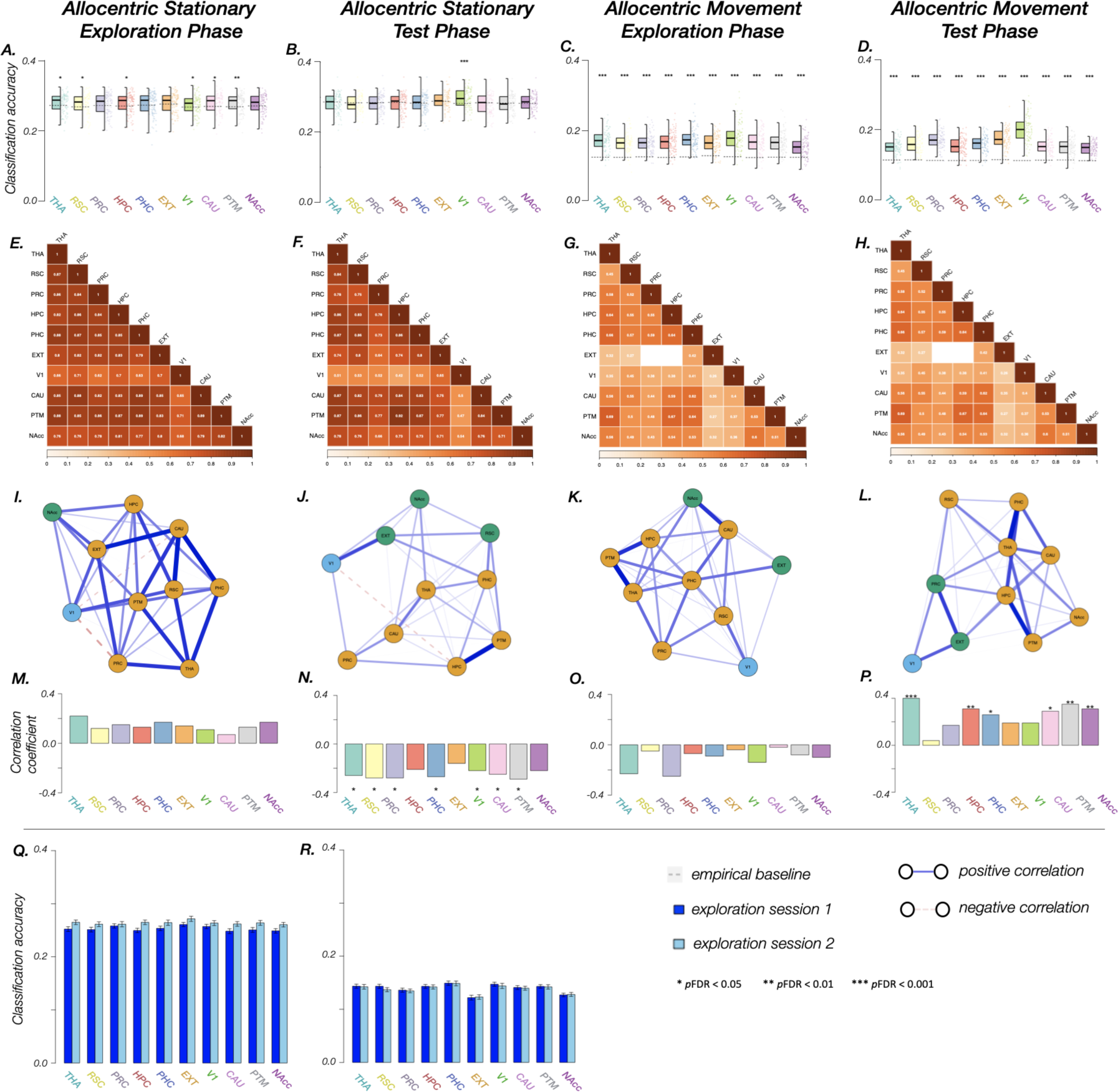
Performance of present allocentric coding (coding the current head direction at the present time) while stationary and present allocentric coding during movement, in 10 ROIs (THA, thalamus; RSC, retrosplenial cortex; PRC, precuneus; HPC, hippocampus; PHC, parahippocampal cortex; EXT, extrastriate cortex; V1, early visual cortex; CAU, caudate; PTM, putamen; NAcc, nucleus accumbens). A.-D. Classification accuracy in each ROI during the four sections of the task: A. allocentric stationary during the exploration phase, B. allocentric stationary during the test phase, C. allocentric movement during the exploration phase, D. allocentric movement during the test phase. E.-H. Pearson correlation matrices of classification accuracy in each pair of ROIs in the four sections of the task. Blank cells indicate non-significant correlations. I.-L. Correlation networks among ROIs in the four sections of the task. Each color represents a separate subnetwork. Note: for better visualization, network connections were tuned with Gaussian Markov random field estimation using graphical LASSO and extended Bayesian information criterion to select optimal regularization parameters. As a result, weak Pearson correlations in correlation matrices may appear as negative connections in the network visualization. Nodes represent ROIs and edges represent tuned correlation coefficients. Subnetworks were generated based on hierarchical clustering and the optimal number of clusters were determined with a silhouette score. M.-P. Correlation coefficients between navigation performance (i.e., accuracy) and classification accuracy in each ROI in the four sections of the task. Q. Significant increase in allocentric stationary classification accuracy between the two exploration sessions (p = 0.043*). R. No change in allocentric classification accuracy between the two exploration sessions during movement.

We also examined periods of visual movement when the paradigm moved the person to the next choice point in the maze. For both the exploration and test phases, during movement, allocentric head direction could be decoded based on patterns of activity within all of our ROIs (Figure 2c, 2d). Interestingly, although classification was similarly successful in many of the brain regions, the correlations between them were much lower than for the stationary period (exploration: r [0.27, 0.69]; test: r [0.26, 0.69]); several smaller sub-networks emerged and both early visual cortex and extrastriate cortex had much lower correlations with all other ROIs. For example, during exploration, retrosplenial cortex was found to be a primary connector between early visual cortex and the rest of the brain, whereas during test extrastriate and precuneus served as the links (Figure 2g, 2h, 2k, 2l). We also found strong correlations between classifier accuracies in the hippocampus, putamen, and thalamus (Figure 2k, 2l), which remained throughout much of our analyses.

### Egocentric Movement Direction Classifiers

In addition to allocentric (cardinal) directions, we also examined whether multiple reference frames could be represented at the same time. To do so, we tested egocentric (left, right, straight) movement directions during the periods of visual movements – the same time points we used for the allocentric movement classification. Note that egocentric stationary only has one direction (straight) and so was not classified. Egocentric movement could be successfully decoded in all of our ROIs during the exploration phase (Figure 3a), and in all except thalamus and extrastriate cortex during the test phase (Figure 3b). Notably, there were no significant correlations in classifier accuracies between extrastriate cortex and the other ROIs during exploration, although the remainder of the correlations were moderately strong (r [0.40, 0.71]; Figure 3c). During the test phase the correlations were very strong, but early visual cortex was only weakly correlated with precuneus (r [0.27, 0.93]; Figure 3d). The network visualizations indicate a shift from a highly interconnected network during exploration (with the exception of extrastriate cortex; Figure 3e) to a sparse network with paired clusters during the test phase (Figure 3f). Together, these findings indicate that both egocentric and allocentric frames of reference are represented in the brain at the same time, with distinct network relationships that are shaped during exploration.

**Figure 3.**
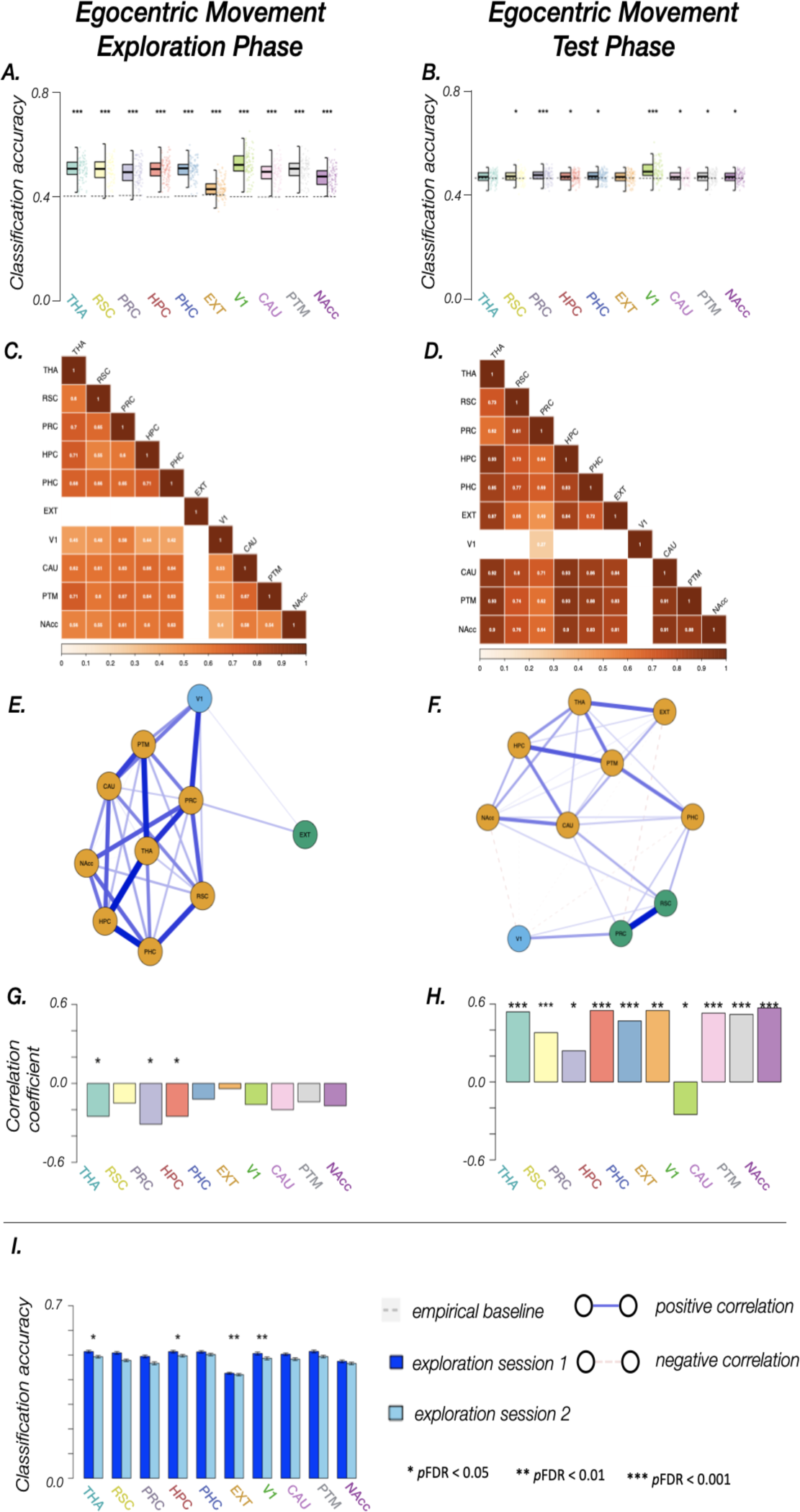
Performance of present egocentric movement in 10 ROIs (THA, thalamus; RSC, retrosplenial cortex; PRC, precuneus; HPC, hippocampus; PHC, parahippocampal cortex; EXT, extrastriate cortex; V1, early visual cortex; CAU, caudate; PTM, putamen; NAcc, nucleus accumbens). A. Classification accuracy in each ROI during egocentric movement during the exploration phase and B. Egocentric movement during the test phase. C.-D. Pearson correlation matrices of classification accuracy in each pair of ROIs in exploration and test phases. Blank cells indicate non-significant correlations. E.-F. Correlation networks among ROIs in exploration and test phases. Each color represents a separate subnetwork. Note: for better visualization, network connections were tuned with Gaussian Markov random field estimation using graphical LASSO and extended Bayesian information criterion to select optimal regularization parameters. As a result, weak Pearson correlations in correlation matrices may appear as negative connections in network visualization. Nodes represent ROIs and edges represent tuned correlation coefficients. Subnetworks were generated based on hierarchical clustering and the optimal number of clusters were determined with silhouette score. G.-H. Correlation coefficients between navigation performance (i.e., accuracy) and classification accuracy in each ROI in the exploration and test phases. I. Significant decrease in egocentric movement classification accuracy between the two exploration sessions (p<0.001***).

### Changes During Learning

We were particularly interested in the dynamics of how these neural representations changed during exploration. Because the exploration was divided into two 8-minute sessions, we compared the classification accuracy across the two sessions in all ROIs. We found a significant main effect of session for allocentric stationary periods (F(1, 95) = 4.13, *p* = .043, η_p_^2^ = .042), with classification strength increasing during exploration (Figure 2q), although there was no difference for the allocentric movement periods (F(1, 95) = .17, *p* = .677, η_p_^2^ = .002) (Figure 2r). Interestingly, classification accuracy for egocentric movement significantly *decreased* both as a main effect (F(1, 95) = 13.71, *p* < .001, η_p_^2^ = .126) and for several ROIs considered individually (i.e., thalamus, putamen, retrosplenial cortex, and precuneus; *ps < 0.05*) (Figure 3i). Taken together, these findings suggest that the representation for allocentric directions strengthened during the course of exploration. In contrast, those for egocentric directions got weaker.

### Classification of Past and Future Trajectories

The previous analyses allowed us to elucidate neural signals about participants’ *present* moment. Critically, they were engaged in a dynamic navigation task that could also provide rich information about their *past and future trajectories*. In addition to processing their current head direction at an intersection, participants were also deciding their path to learn the maze during exploration, as well as planning trajectories towards the target goal during the test phase. To examine future trajectory information within our ROIs, we attempted to decode both an individual’s future (one step ahead) and past (one step behind, see Supplement) movement using the same time points and data we used for the allocentric stationary classifiers. Intriguingly, allocentric future movements could be decoded during exploration within all ROIs except the nucleus accumbens (Figure 4a). During the test phase, we could successfully classify future allocentric movement in thalamus, putamen, parahippocampal cortex, and early visual cortex (Figure 4b). Classification for almost all regions in the network were highly correlated with each other during exploration, although with more moderate correlations with V1 (r [0.70, 0.96]; Figure 4e, 4i). The network was also fairly highly correlated during the test phase (r [0.34, 0.90]), but now two large sub-networks emerged, both of which spanned visual, medial temporal, and basal ganglia regions (Figure 4f, 4j). A qualitatively similar pattern of results was seen for past allocentric movements, with all regions successfully classifying during exploration, although no regions passed significance during test (Figure S2).

**Figure 4.**
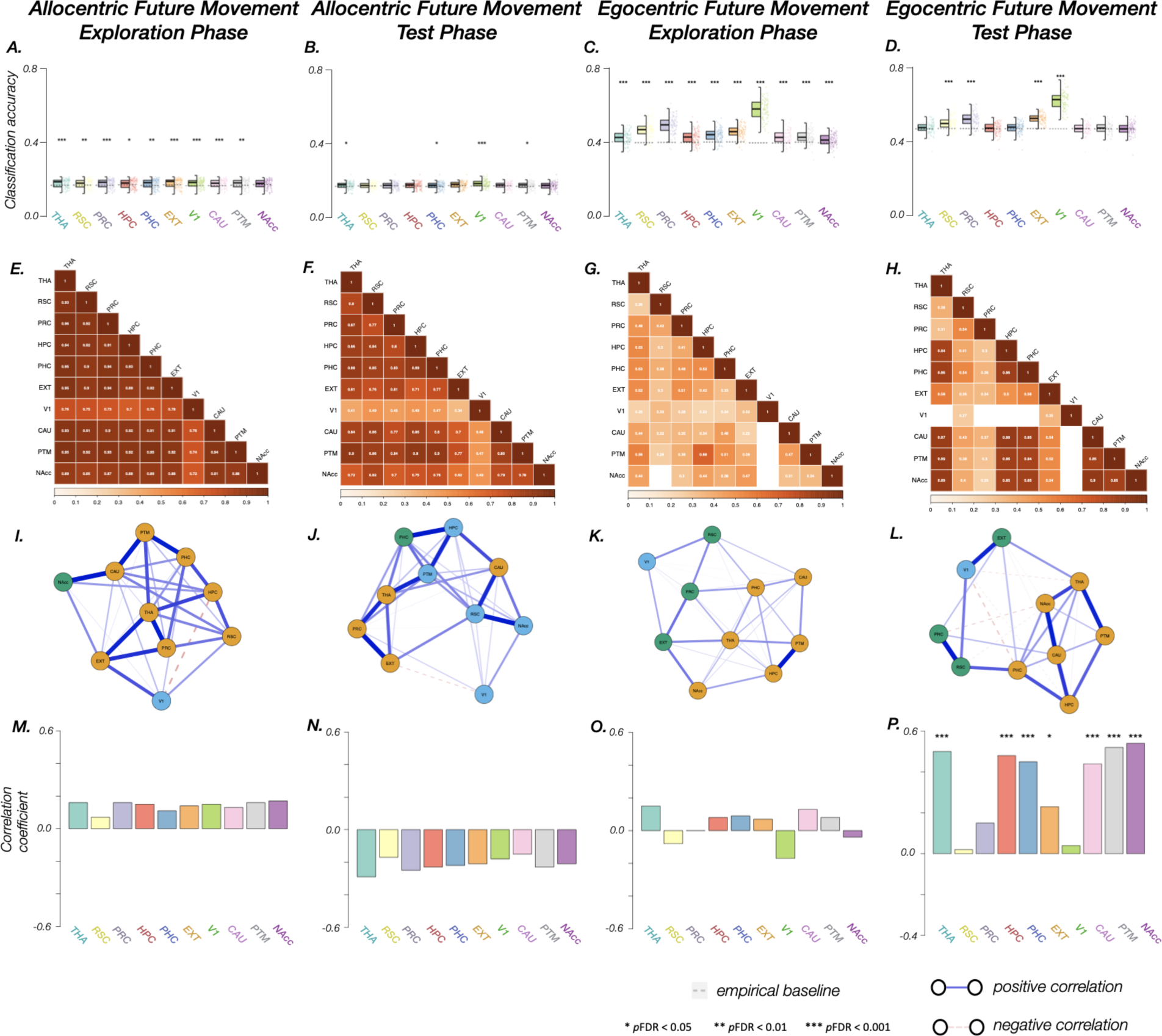
Performance of allocentric and egocentric future movement in 10 ROIs (THA, thalamus; RSC, retrosplenial cortex; PRC, precuneus; HPC, hippocampus; PHC, parahippocampal cortex; EXT, extrastriate cortex; V1, early visual cortex; CAU, caudate; PTM, putamen; NAcc, nucleus accumbens). These classifications are when the participant is stationary at an intersection, but we classified based on their next allocentric or egocentric movement. A.-D. Classification accuracy in each ROI during the four sections of the task: A. allocentric future movement during the exploration phase, B. allocentric future movement during the test phase, C. egocentric future movement during the exploration phase, C. egocentric future movement during the test phase. E.-H. Pearson correlation matrices of classification accuracy in each pair of ROIs in the four sections of the task. Blank cells indicate non-significant correlations. I.-L. Correlation networks among ROIs in the four sections of the task. Each color represents a separate subnetwork. Note: for better visualization, network connections were tuned with Gaussian Markov random field estimation using graphical LASSO and extended Bayesian information criterion to select optimal regularization parameters. As a result, weak Pearson correlations in correlation matrices may appear as negative connections in network visualization. Nodes represent ROIs and edges represent tuned correlation coefficients. Subnetworks were generated based on hierarchical clustering and the optimal number of clusters were determined with silhouette score. M.-P. Correlation coefficients between navigation performance (i.e., accuracy) and classification accuracy in each ROI in the four sections of the task.

Egocentric future movements were also significantly classified in all regions during exploration (Figure 4c). During the test, we were able to successfully classify future egocentric movements in the retrosplenial cortex, precuneus, extrastriate cortex, and early visual cortex (Figure 4d). Very similar results were seen for past egocentric movements (Figure S2). The correlations between egocentric classifications during exploration demonstrated a weakly but fairly evenly connected network (r [0.22, 0.68]; Figure 4g, 4k) with one relatively strong correlation between classifiers in hippocampus and putamen. During test the network analysis revealed several sub-networks, with early visual cortex only correlating with retrosplenial and extrastriate cortices and highly variable correlations between all ROIs (r [0.25, 0.90]; Figure 4h, 4l). Retrosplenial cortex appeared to serve as a bridge to the strong subnetwork of the remaining ROIs via the parahippocampal cortex. Thus, not only present *but future and past information* about the travel trajectory was represented in these brain areas during active navigation.

### Relationship with Navigation Performance

We sought to determine whether the strength of the classification accuracy was associated with subsequent performance in the navigation test. During the exploration phase, none of the egocentric or allocentric classifiers (including past, present, and future) showed any positive relationships between classification accuracy and behavioral accuracy on the task (Figure 2m, 2o, 4m, 4o, S2), with a few egocentric classifiers showing significant negative correlations (Figures 3g). This result suggests that the representations we found during exploration are broad signals, largely common across all individuals regardless of their eventual ability to learn the path layout, which is known as the graph structure of the environment^27–30^. We only found one meaningful correlation during exploration with eventual path efficiency (Figure S3a), for allocentric representations while stationary in thalamus. This result suggests that allocentric representations that were being built in thalamus during the exploration process eventually were translated into metric knowledge of the environment, for good learners.

In contrast, in the test phase we found a large number of relationships between classification strength and performance in the task. Surprisingly, we found a negative relationship between classification strength of the stationary allocentric head direction classifier and both accuracy and path efficiency for almost all ROIs (r [-0.21, −0.29]; Figure 2n, Figure S3b), as well as for past allocentric trajectories (r [-0.23, −0.29]; Figure S2n). During movement in the test phase, there was a positive relationship between allocentric classifier strength and performance in thalamus, hippocampus, parahippocampal cortex, caudate, putamen, and nucleus accumbens (r [0.26, 0.40]) (and with path efficiency in thalamus, hippocampus, caudate, putamen, and nucleus accumbens), such that people with stronger classification strength in those areas during test also performed better in the task (Figure 2p, Figure S3d).

More strikingly, during the test phase the relationship between egocentric classification strength and performance in the navigation task was very strong and positive throughout most of our ROIs. For egocentric movement during the test phase, classification strength was positively correlated with accuracy (r [0.24, 0.57]) (with similar relationships in nearly all ROIs for path efficiency) in all regions, except for early visual cortex, which was significantly negatively correlated (r = −0.25) (Figure 3h, Figure S3f). When stationary, performance was remarkably correlated with the classification strength of future egocentric movements in thalamus, hippocampus, parahippocampal cortex, extrastriate cortex, caudate, putamen, and nucleus accumbens (r [0.23, 0.54]) (with most of these same areas having a relationship with path efficiency) (Figure 4p, S3j). Past egocentric movement classification was correlated with accuracy in all ROIs except retrosplenial cortex (r [0.26, 0.57]; Figure S2p), with similar findings for path efficiency (Figure S3n).

Together, these findings suggest that while processing movement, both allocentric and egocentric information is useful for performance. However, when stationary and planning movement trajectories, better navigators have converted their existing path knowledge into egocentric signals.

## Discussion

Using multivariate pattern classification paired with an fMRI navigation task, we found evidence of simultaneous tuning to multiple features of head and travel direction in a distributed network in the human brain. This network 1) had strong connections with both allocentric and egocentric movement trajectories, 2) changed during the course of exploration, 3) coded for past and future movements as well as the present head direction, and 4) had classification strengths that were strongest for individuals who translated their future trajectories into egocentric movements once they learned the environment. Our findings provide detailed insight into the dynamics of trajectory planning in humans and move well beyond existing studies that have looked for evidence of individual brain structures performing individual aspects of navigation. These results support a richer framework of navigation, which posits that multiple computations are dynamically performed simultaneously.

Our findings extend previous findings in humans and rodents demonstrating that thalamus, retrosplenial cortex, early visual cortex, and precuneus support head and travel direction^1,3,12–15,17,25^. Our results demonstrate a wider network of regions that support both egocentric and allocentric travel direction, including hippocampal and parahippocampal cortices, basal ganglia, and extrastriate visual regions. We found multiple sub-networks that support distinct aspects of head and travel information and that differ across exploration and test. In particular, we frequently observed correlations between classifier strengths in retrosplenial cortex, early visual cortex, extrastriate cortex, and precuneus as well as correlations between hippocampus, thalamus, and putamen.

Beyond the direct classification of present heading or travel direction, we also found evidence of past and future trajectory coding at the same time points that we coded for allocentric direction. These findings suggest that these networks could be important for goal planning and decision making^31–34^. The rodent literature has found trajectory mapping in retrosplenial cortex^9,10^, and hippocampal coding of both spatial and non-spatial trajectories^4–6,35^. Our findings provide important insight into how these trajectory mappings occur in the human brain.

We also found a dynamic relationship between classifier strengths and performance in the task. During exploration, there were no significant positive and some significant negative relationships between classifier strengths and eventual accuracy on the navigation task. During the test period, the stationary allocentric classifiers had significant negative correlations with task performance. In contrast, the exact same stationary time points had significant positive correlations with performance when classified based on the previous (past) and the following (future) egocentric movement. When presently moving, both allocentric and egocentric classifiers in the test phase were correlated with performance. These results suggest that better navigators convert their path knowledge into egocentric signals during planning, and can rely on both egocentric and allocentric signals when moving. Notably, the hippocampus was among the regions where classifier strength of egocentric movements was associated with better performance. These findings challenge the standard perspective that the hippocampus is largely related to allocentric coding, and instead suggest that it is also important for egocentric processing. Furthermore, these findings challenge the dominant framework that allocentric coding is the primary mechanism for organisms to successfully navigate through complex environments.

Together, the findings of this study call for a new framework of human navigation. We propose that the brain computes multiple co-existing representations of both egocentric and allocentric reference frames across a network of brain regions. Egocentric and allocentric reference frames trade off during learning, and egocentric codings may ultimately be most useful for reaching navigational goals if the navigator knows the environment. These multiple computations are not limited to the present time, but extend into the future and connect to the past, with the memory of previous trajectories influencing travel planning.

Although we see strong evidence of coding of head and travel direction signals in the visual cortex, it is unlikely that the visual information alone is driving our results. Indeed, classification of the present allocentric stationary condition had generally poor accuracy during the test phase - except in early visual cortex - despite this being a fairly straightforward time to use visual information to determine head direction. It appears that other representations, such as egocentric and future planning, are becoming stronger, thereby minimizing the effects of simple visual features on our classification.

We acknowledge that the navigation trajectories in our study were purely based on visual information; full proprioceptive and vestibular information could alter the nature of these head direction representations^36^. fMRI classification methods cannot always distinguish between independent coding in a brain region and strong connections from an input area that codes for that information. Likewise, declines in classifier accuracy during learning could reflect direct changes in that region’s processing or could indicate multiple sensitivities that average out. Combining human imaging work with current and future animal studies examining head direction and egocentric and allocentric representations can help disentangle these factors.

By examining current, past, and future trajectory planning, our findings provide evidence of the dynamic nature of the brain networks that support active and complex navigation in humans and provide insights into the nature of how individuals acquire and use spatial knowledge to plan trajectories in complex environments. Together, these findings mark a foundational shift away from the dominant framework of allocentric coding and instead support a new framework of navigation that includes multiple co-existing dynamic computations.

## Acknowledgments

The data for this study was collected at the University of California, Santa Barbara Brain Imaging Center. The authors would like to thank Rie Davis, Grace Nicora, Justin Kasowski, Rob Woodry, Alina Tu, Andrew Huang, Mathias Goncalves, Mario Mendoza, Scott Grafton, Emily Grossman, Bruce McNaughton, Jeff Krichmar, Ramesh Srinivasan, and Craig Stark.

## Author Contributions

**Y.C.** methodology, investigation, formal analysis, conceptualization, data curation, writing - original draft, visualization; **S.L.** conceptualization, writing - review and editing, funding acquisition; **C.S.** conceptualization, writing - review and editing, funding acquisition; **E.C.** conceptualization, methodology, writing - original draft, writing - review and editing, supervision, project administration, funding acquisition

## Materials and Methods

### Key resources table

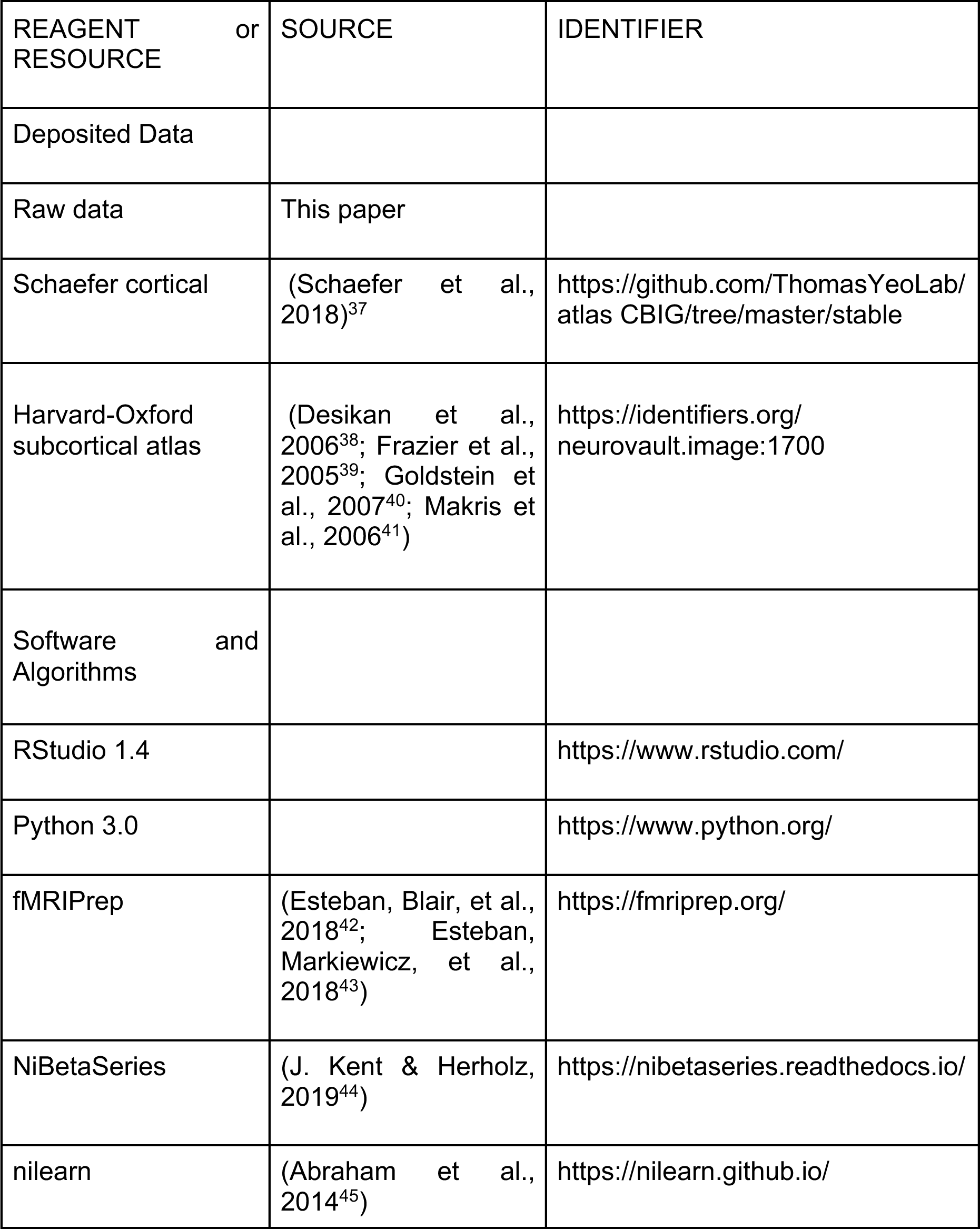
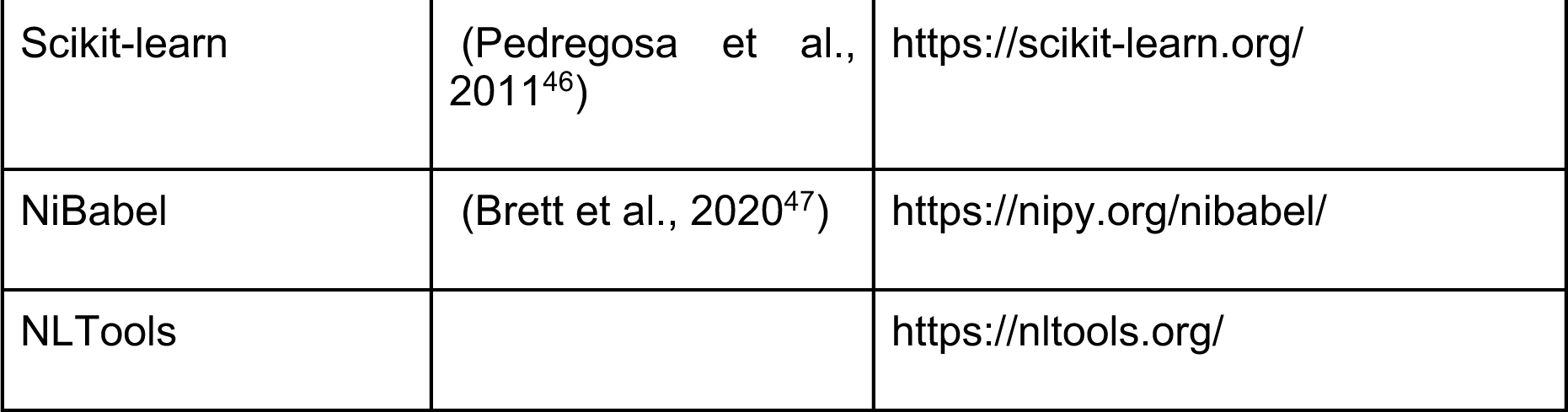

### Participants

113 participants were recruited through previous studies, campus flyers, or word of mouth. Participants were pre-screened for neurological disease, psychotropic medication, and MRI safety compliance. Fifteen of those recruited failed to complete the experiment. Of these, five participants failed to schedule/show up the scan portion of the experiment, one subject was discarded due to technical issues, and nine subjects could not complete the study due to motion sickness or problems being in the MRI machine (e.g., claustrophobia). Further, we excluded two participants with high motion in the exploration phase, and seven participants with high motion in the test phase. Thus, the final dataset consisted of 96 participants (44 females, 52 males; age 18-37, mean = 20.75, s.d = 3.17) for the exploration phase, and 91 participants (44 females, 47 males; age 18-37, mean = 20.79, s.d = 3.19) for the test phase (all 91 participants in the test phase dataset were also included in the exploration phase dataset).

### Virtual environments and navigation interface

The desktop virtual maze environment (Figure 1) was adapted from tasks the lab has used previously^19,27,30,48^ and was designed to produce a wide range of performance. The environment consisted of several main hallways with branch alcoves containing nine target objects. Paintings on the walls of hallways served as landmarks to aid in orientation. Participants pressed arrow keys to move around the environment, with translations fixed at 1.0 m/s (virtual meters/second) and rotation speed fixed at 90◦ per second. Movement was gated such that the button presses at each choice point caused visual movement to the next choice point. This was achieved by recording videos of the movement between locations. The correct video would play based on the person’s location, facing direction, and the choice they made (left turn, right turn, or straight). A static image of each choice point was created by taking the first image of each video and adding the arrows of the potential choices in Photoshop (Adobe). The maze was created in Blender and rendered in Unity. The lab recorded videos from Unity and presented the experiment using E-Prime (Psychology Software Tools) software.

### Behavioral task

The Maze Learning Task was designed to assess the ability to learn the graph structure of an environment—the paths and connections between locations^19,27,30^. A video example of a test trial is available online at https://www.youtube.com/watch?v=LMsGpo2Ss7M.

#### Exploration phase

Participants navigated the virtual environment for 16 minutes, during two 8-minute fMRI scan runs, using a button box to make right or left turns or straight movement (Figure 1b). All participants were instructed to find all of the objects and learn their locations. Four different start locations were used; the start location was counterbalanced for each participant for each of the two runs. The exploration phase was the primary time to learn the environment, although some learning was possible during the test.

#### Test phase

Acquired spatial knowledge was tested on each trial by starting participants at one object in the maze, and then directing them to travel, via the shortest route, to another object using the hallways of the maze. Participants then used the button box to move through the maze like they did in the exploration phase, pressing a button when they thought they reached the target object. Feedback was minimized by changing all of the objects in the maze (including the start and target objects) into red spheres during the test phase (Figure 1c), although the landmark paintings in the hallway remained. The participants could be instructed either to “Start at the spaceship, go to the clock”, or to “Go to the clock, start at the spaceship” and were informed of the subtle differences in wording prior to starting the trials. The start and target objects were each displayed for between 3-5 seconds, with the specific timing drawn from a random distribution in the range. This task tests graph knowledge^27,30^ because it requires knowing the connections of the hallways to reach the target without necessarily knowing the metric distances and angles between locations. Trials had a 45 second time limit. When the participant made a selection or when time expired, a 6-second (with random jitter) inter-trial interval began.

#### Procedure

One or two days prior to imaging, participants were greeted in the lab and were given information about the study. They signed consent forms indicating their consent to participate in the study and completed several paper and pencil spatial abilities tasks.

The following day at the scanner, participants were given instructions about how to use the controller and given a short practice in a different environment to show them how the movement worked. They were instructed to find all the objects and learn their locations.

Participants changed into scrubs and were shown the scanning equipment. After aligning the participant in the scanner and ensuring their comfort, including giving them earplugs and cushions, the scan session began. The first set of anatomical scans was collected (see Image Acquisition), then the experimental task began.

Participants completed two 8-minute scan runs where they were free to explore the maze. At the end of the first run, participants were informed of how many objects they located and how many were left undiscovered.

After the exploration phase, participants were given instructions for the test trials. They completed 6 scan runs consisting of 8 trials each (total = 48 trials). With nine objects in the maze, a full permutation of all start and target pairs yielded 72 possible trials. However, due to time constraints, a subset of 48 trials was used for each participant. Three lists of trials were used, each with 2/3 of the possible trials. Each participant followed one of the lists, counterbalanced across participants. The trial list was presented in random order for each participant.

### Navigation metrics

#### Accuracy

Participants were scored by their proportion of correct trials. Trials were considered “correct” if the participant ended at the target object, regardless of whether they pressed the “selection” button or whether they ran out of time when just reaching the target. Proportion correct scores ranged from around chance performance (chance is defined as .111, or ending at 1 out of the 9 possible target objects) to perfect. Only 4 participants were correct on every trial.

#### Path efficiency

Participants were scored by their average path efficiency across trials. For each trial, the path efficiency score is path distance traveled (by the participant) divided by shortest path distance between the start and target location. We term this measure excess path ratio. Therefore, the excess path ratio ranges between 1 to infinity, where a score of 1 means the participant took the most efficient path possible and high ratios indicate very inefficient paths.

#### Allocentric stationary facing directions

Based on each participant’s individual behavioral travel trajectory, we constructed design matrices for each participant’s stationary facing directions. There were 4 stationary facing directions when standing still at an intersection: north, south, east, or west. Notice that the directions were arbitrarily defined, but different labels should not affect our analyses. Because participants were freely moving in the maze in both the exploration and test phases, we did not expect any two participants would share the same trajectory. Therefore, we constructed one design matrix for the exploration phase and one design matrix for the test phase separately for each person.

We did not construct a design matrix for egocentric stationary facing directions because there was only one egocentric stationary facing direction (facing forward).

#### Allocentric movements

Based on each participant’s individual behavioral travel trajectory, we constructed design matrices for each participant’s present allocentric movements. There were 12 allocentric present movements: moving straight north, south, east, west, or turning from north to east, east to south, south to west, west to north, north to west, west to south, south to east, and east to north. More specifically, we constructed one design matrix for the exploration phase and one design matrix for the test phase separately for each person.

Because the maze environment was largely grid-based, almost all of the turns were 90◦, so we did not include the very few 180◦ turning events (i.e., south to north, east to west, typically at dead ends) in the analyses.

#### Egocentric movements

Based on each participant’s individual behavioral travel trajectory, we constructed design matrices for each participant’s present egocentric movements. There were 3 egocentric movements: forward/straight, turning left (counterclockwise) and turning right (clockwise). More specifically, we constructed one design matrix for the exploration phase and one design matrix for the test phase separately for each person.

#### Allocentric future and past movements

Based on each participant’s individual behavioral travel trajectory, we constructed design matrices for each participant’s allocentric future movements (i.e., the next move that they made, using an allocentric reference frame) when they were stationary at an intersection deciding where to go next. There were 12 allocentric future movements: moving straight north, south, east, west, or turning from north to east, east to south, south to west, west to north, north to west, west to south, south to east, and east to north. More specifically, we constructed one design matrix for the exploration phase and one design matrix for the test phase separately for each person.

Allocentric past movements were constructed the same way, but using the previous move that the participant had made, using an allocentric reference frame.

#### Egocentric future and past movements

Based on each participant’s individual behavioral travel trajectory, we constructed design matrices for each participant’s egocentric future movements (i.e., the next move that they made, using an egocentric reference frame) when subjects were stationary at an intersection deciding where to go next. There were 3 egocentric future movements: moving forward, turning left (counterclockwise), or turning right (clockwise). More specifically, we constructed one design matrix for the exploration phase and one design matrix for the test phase separately for each person.

Egocentric past movements were constructed the same way, but using the previous move that the participant had made, using an egocentric reference frame.

### fMRI acquisition

MRI imaging was conducted on a 3.0T Siemens Prisma magnetic resonance imaging system at the UC Santa Barbara Brain Imaging Center using a 64-channel head coil. At the beginning of the scan session, approximately twenty minutes of anatomical scans were acquired, including a magnetization-prepared, rapid-acquisition gradient-echo (MPRAGE) T1 weighted sequence image (TR = 2500ms, TE = 2.2ms, 7° flip angle, FOV 256mm × 256mm, voxel size 0.9mm × 0.9mm × 0.9mm), field maps, and one scan of diffusion weighted images. A 7-minute task-free resting state functional scan was also acquired. Following this, two 8-minute task-based functional scans of the exploration phase of the maze task and six sets of task-based functional scans were acquired for the testing phase of the maze task; this section lasted approximately 40-50 minutes (TR = 720ms, TE = 37ms, 52° flip angle, FOV 208mm x 208mm, voxel size 2.0mm x 2.0mm x 2.0mm, 72 slices, multi-band acceleration = 8). After the task, another twenty minutes of anatomical scans were collected, including a second set of diffusion weighted images, and several T2 weighted images. The functional images analyzed here were obtained as part of the larger study consisting of both fMRI and anatomical images; the anatomical portion (i.e., diffusion images and T2) and resting state data are not analyzed or reported here.

### fMRI preprocessing

Results included in this manuscript come from preprocessing performed using fMRIPrep 1.5.10 (RRID:SCR 016216^42,43^), which is based on Nipype 1.4.2 (RRID:SCR 002502^49,50^).

### Anatomical data preprocessing

The T1-weighted (T1w) image was corrected for intensity non-uniformity (INU) with N4BiasFieldCorrection^51^, distributed with ANTs 2.2.0 (RRID:SCR 004757^52^), and used as T1w-reference throughout the workflow. The T1w-reference was then skull-stripped with a Nipype implementation of the antsBrainExtraction.sh workflow (from ANTs), using OASIS30ANTs as target template. Brain tissue segmentation of cerebrospinal fluid (CSF), white-matter (WM) and gray-matter (GM) was performed on the brain-extracted T1w using fast (FSL 5.0.9, RRID:SCR 002823^53^). Brain surfaces were re-constructed using recon-all (FreeSurfer 6.0.1, RRID:SCR 001847^54^), and the brain mask estimated previously was refined with a custom variation of the method to reconcile ANTs-derived and FreeSurfer-derived segmentations of the cortical gray matter of Mindboggle (RRID:SCR 002438^55^). Volume-based spatial normalization to one standard space (MNI152NLin2009cAsym) was performed through nonlinear registration with antsRegistration (ANTs 2.2.0), using brain-extracted versions of both T1w reference and the T1w template. The following template was selected for spatial normalization: ICBM 152 Nonlinear Asymmetrical template version 2009c (RRID:SCR 008796^56^, TemplateFlow ID:MNI152NLin2009cAsym).

### Functional data preprocessing

For each of the 8 BOLD runs found per subject (across the two exploration session scan runs and the 6 test trial runs), the following preprocessing was performed. First, a reference volume and its skull-stripped version were generated using a custom methodology of fMRIPrep. A B0-nonuniformity map (or fieldmap) was estimated based on a phase-difference map calculated with a dual-echo GRE (gradient-recall echo) sequence, processed with a custom workflow of SDCFlows inspired by the epidewarp.fsl script and further improvements in HCP Pipelines^57^. The fieldmap was then co-registered to the target EPI (echo-planar imaging) reference run and converted to a displacements field map (amenable to registration tools such as ANTs) with FSL’s fugue and other SDCflows tools. Based on the estimated susceptibility distortion, a corrected EPI (echo-planar imaging) reference was calculated for a more accurate co-registration with the anatomical reference. The BOLD reference was then co-registered to the T1w reference using bbregister (FreeSurfer) which implements boundary-based registration^58^. Co-registration was configured with six degrees of freedom. Head-motion parameters with respect to the BOLD reference (transformation matrices, and six corresponding rotation and translation parameters) are estimated before any spatiotemporal filtering using mcflirt (FSL 5.0.9^59^). The BOLD time-series (including slice-timing correction when applied) were resampled onto their original, native space by applying the transforms to correct for head-motion. These resampled BOLD time-series will be referred to as preprocessed BOLD in original space, or just preprocessed BOLD. BOLD runs were slice-time corrected using 3dTshift from AFNI 20160207^60^ (RRID:SCR 005927). The BOLD time series were resampled to surfaces on the following spaces: fsaverage5. The BOLD time-series (including slice-timing correction when applied) were resampled onto their original, native space by applying a single, composite transform to correct for head-motion and susceptibility distortions. These resampled BOLD time-series will be referred to as preprocessed BOLD in original space, or just preprocessed BOLD. The BOLD time-series were resampled into standard space, generating a preprocessed BOLD run in MNI152NLin2009cAsym space. First, a reference volume and its skull-stripped version were generated using a custom methodology of fMRIPrep. Several confounding time-series were calculated based on the preprocessed BOLD: framewise displacement (FD), DVARS and three region-wise global signals. FD and DVARS were calculated for each functional run, both using their implementations in Nipype (following the definitions by^61^). The three global signals were extracted within the CSF, the WM, and the whole-brain masks. Additionally, a set of physiological regressors were extracted to allow for component-based noise correction (CompCor^62^). Principal components were estimated after high-pass filtering the preprocessed BOLD time-series (using a discrete cosine filter with 128s cut-off) for the two CompCor variants: temporal (tCompCor) and anatomical (aCompCor). tCompCor components were then calculated from the top 5% variable voxels within a mask covering the subcortical regions. This subcortical mask is obtained by heavily eroding the brain mask, which ensures it does not include cortical GM regions. For aCompCor, components were calculated within the intersection of the aforementioned mask and the union of CSF and WM masks calculated in T1w space, after their projection to the native space of each functional run (using the inverse BOLD-to-T1w transformation). Components were also calculated separately within the WM and CSF masks. For each CompCor decomposition, the k components with the largest singular values were retained, such that the retained components’ time series are sufficient to explain 50 percent of variance across the nuisance mask (CSF, WM, combined, or temporal). The remaining components were dropped from consideration. The head-motion estimates calculated in the correction step were also placed within the corresponding confounds file. The confound time series derived from head motion estimates and global signals were expanded with the inclusion of temporal derivatives and quadratic terms for each^63^. Frames that exceeded a threshold of 0.5 mm FD or 1.5 standardized DVARS were annotated as motion outliers. All resamplings can be performed with a single interpolation step by composing all the pertinent transformations (i.e. head-motion transform matrices, susceptibility distortion correction when available, and co-registrations to anatomical and output spaces). Gridded (volumetric) resamplings were performed using antsApplyTransforms (ANTs), configured with Lanczos interpolation to minimize the smoothing effects of other kernels^64^. Non-gridded (surface) resamplings were performed using mri vol2surf (FreeSurfer).

Many internal operations of fMRIPrep use Nilearn 0.6.2 (RRID:SCR 001362^45^), mostly within the functional processing work-flow. For more details of the pipeline, see the section corresponding to workflows in fMRIPrep’s documentation.

### Beta series analysis

First level analyses were conducted using the beta series analysis method^65^, which has been used for previous navigation studies^16,66^. The beta series method utilizes the univariate fMRI data analysis so that parameter estimates (i.e., beta weights), reflecting the magnitude of the task-related blood oxygen level dependent (BOLD) responses are estimated for each trial. Therefore, the beta series analysis requires that the individual trials of events examined in the analysis be modeled separately.

It is worth noting that, because all participants were allowed to freely move in the virtual environment during both exploration and test phases, the type of behavior (moving or stationary, the particular facing direction) at different time points will be different. Therefore, we did not create a design matrix before the experiment, but constructed a design matrix for each trial of each participant separately based on their behavioral data. The individualized design matrices were then implemented for first level analyses.

Results included in this manuscript come from modeling performed using NiBetaSeries 0.6.0^44^, which is based on Nipype 1.4.2^49,50^.

### Beta series modeling

In addition to condition regressors, csf, white matter, global signal, trans x, trans y, trans z, rot x, rot y, rot z, framewise displacement, motion outlier* and a high-pass filter of 0.0078125 Hz (implemented using a cosine drift model) were included in the model. ‘Global signal’ refers to global signal processing. This confounder was included as suggested by^67^; ‘trans x, trans y, trans z, rot x, rot y, rot z’ refers to six degrees of head motion. Framewise displacement is a measurement of head motion from one voxel to the next. This confounder was included as it was shown to improve performance in groupwise analyses^68^; Motion outlier is a type of volume censoring that discards problematic time points (* refers to wildcard matching of all motion outliers for each subject). These confounders were included as it was shown to be effective in reducing motion-related artifacts^69^. AR(1) prewhitening was applied in each model to account for temporal autocorrelation.

After fitting the model, the parameter estimate (i.e., beta) map associated with the target trial’s regressor was retained and concatenated into a 4D image with all other trials from the same condition, resulting in a set of N 4D images, where N refers to the number of conditions in the task. The number of volumes in each 4D image represents the number of trials in that condition. The same condition here means the same type of event. For example, in generating the betaseries for stationary allocentric facing directions (north, east, south, west), all “north” event-related images were concatenated as one 4D image for each run in each subject.

### Software Dependencies

Additional libraries used in the NiBetaSeries workflow include Pybids 0.9.5^70^, Niworkflows 1.0.4, Nibabel 3.1.0, Pandas 0.24.2^71^, and Numpy 1.20.3^72,73^.

### ROI analyses

Our principal fMRI analyses were conducted using targeted a priori regions of interest (ROIs) (see Figure S1). Previous studies strongly suggest task-evoked directional activation in thalamus^12^, retrosplenial cortex^12,13,17,26^, precuneus^12,15^, extrastriate cortex^16^, and early visual cortex^13,17,25^. Because allocentric navigation function is typically centered around the hippocampus^21,26^, we also selected the hippocampus as an ROI. We were also interested in directional encoding for allocentric compared to egocentric frames of reference, and so we included basal ganglia regions, which are important for navigation with an egocentric strategy^22–24^ and with decision making^74–76^ (i.e., caudate, putamen, and nucleus accumbens). We did not include entorhinal cortex because segmenting entorhinal cortex requires a high-resolution image of the medial temporal lobe.

All cortical ROIs (including retrosplenial cortex, precuneus, extrastriate cortex, parahippocampal cortex, and early visual cortex) were generated for each participant’s brain in the 521MNI space based on the Schaefer 100 regions atlas (https://github.com/ThomasYeoLab/CBIG/tree/master/ stable projects/brain parcellation/Schaefer2018 LocalGlobal)^37^.

All subcortical ROIs (including thalamus, hippocampus, caudate, putamen, and nucleus accumbens) were generated for each participant’s brain in the 521MNI space based on the Harvard-Oxford subcortical probabilistic atlas with 25% threshold (https://identifiers.org/neurovault.image:1700)^38–41^. All ROIs included combined bilateral regions.

### Multivariate pattern analysis

To derive an index of trial-wise directional representation in eight different directional behaviors (see Navigation metrics; e.g., north, east, south, west for stationary facing direction), we computed category-selective pattern using multivariate pattern analyses (MVPA). Under the assumption that each person has a unique cognitive map, we conducted MVPA for each participant, with separate exploration (i.e., learning) and test phases. Because we were interested in head direction under both egocentric and allocentric frames of reference, we focused on activity in 10 ROIs (see ROI analyses). MVPA was performed using nilearn (https://nilearn.github.io/), Scikit-learn (https://scikit-learn.org/), NLTools (https://nltools.org/) packages, and Python scripts.

The multi-class classification was performed using a gaussian-kernelized support vector machine (SVM) with nested cross-validation under L2 regularization, based on a one-versus-one classifier. For a better estimate of the generalization performance, we used a 3-fold cross validation to evaluate the combination of two parameters: gaussian kernel width and regularization penalization parameter (see Equation 3 and Equation 2). The data were first shuffled and then randomly split into 3 sets. For each parameter setting (grid search among 6 values (i.e., 0.001, 0.01, 0.1, 1, 10, 100) for each parameter, respectively), 3 accuracy values were computed, one for each split in the cross-validation. Then the parameter setting with the highest mean validation accuracy was chosen. The 3-fold inner cross-validation was nested under a 10-fold outer cross-validation, where the dataset was first shuffled and then randomly split into 10 sets. The decoder was trained on 9 of the sets, and the performance was tested on the final set. This was done 10 times in a rotating fashion, so that each set was tested once. The performance on all test sets was generally averaged together to determine the overall performance (see Equation 1 and Equation 2). The performance was measured by accuracy.

The calculation follows the equations below:

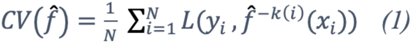

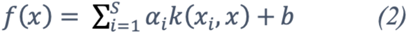

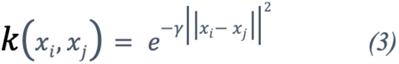

(1) Cross-validation function. *CV* is the accuracy. *N* refers to the number of folds. *K:* {1,…, *N}. L* refers to loss function. *f̂* refers to the predicted label (e.g., north), calculated by equation (2). *y_i_* refers to actual labels (e.g., north). *x_z_* refers to neural signal (i.e., beta signal of each event). *N =* 10.
(2) Kernel SVM function. *x_i_* refers to support vectors, s refers to the number of support vector. *α_i_* refers to coefficient, *b* is the parameter learned in the training phase, *k* is the kernel function, calculated by equation (3).
(3) Gaussian radial basis function. *x_i,_x_j_* are data points, | |*x_i_ - x_j_*| |^2^ denotes Euclidean distance, *γ* is the parameter that controls the width of the Gaussian kernel.

ROI masks were created using the Harvard-Oxford subcortical atlas and the Schaefer cortical atlas (see ROI analyses) and then applied to beta series from the whole brain voxels (see Beta series analysis) for each event type. To improve classification accuracy, we conducted a within-run mean-centering method for trial specific estimates^77^ (i.e., subtracting each voxel’s run-level mean across trials of all types within each run) with the exception of using default scaling for analyses that does not have all event types within each run (the test phase of the allocentric present direction, allocentric past direction, allocentric future direction analyses).

The theoretical baseline was not implemented due to not fully balanced movements in each direction in each participant’s active navigation. Therefore, a more conservative empirical baseline was established using the same analyses pipeline described above with a meta permutation-based test (take the average of all participants’ tests with label shuffles) for each ROI.

### Quantification and Statistical Analyses

A one-sample t-test was conducted for group classification accuracy compared with empirical baseline, for each ROI, at exploration and test phases, respectively (with FDR correction for multiple comparisons^78–80)^. To compare the model classification strength between the two exploration sessions, we first down sampled events in the second exploration phase to be equivalent to the event number in the first exploration session by randomly taking out ∼10 events. Then we conducted session (session 1, session 2) x ROI (10 ROIs) two-way ANOVA analyses, with Tukey posthoc tests to compare the difference between the two exploration sessions. We also conducted ROI clustering based on correlation matrices of ROI classification accuracy in each phase using agglomerative hierarchical clustering^81^. The optimal clusters were determined with silhouette score^82^ and visualized with different colors in networks. We conducted Pearson correlation of classification accuracy between every pair of ROIs, between behavioral performance accuracy and the classification accuracy for each ROI, between path efficiency and the classification accuracy for each ROI, at exploration and test phases, respectively (with FDR correction for multiple comparisons^78–80)^. All statistical analyses were conducted in RStudio.

## Supplement

**Figure S1.**
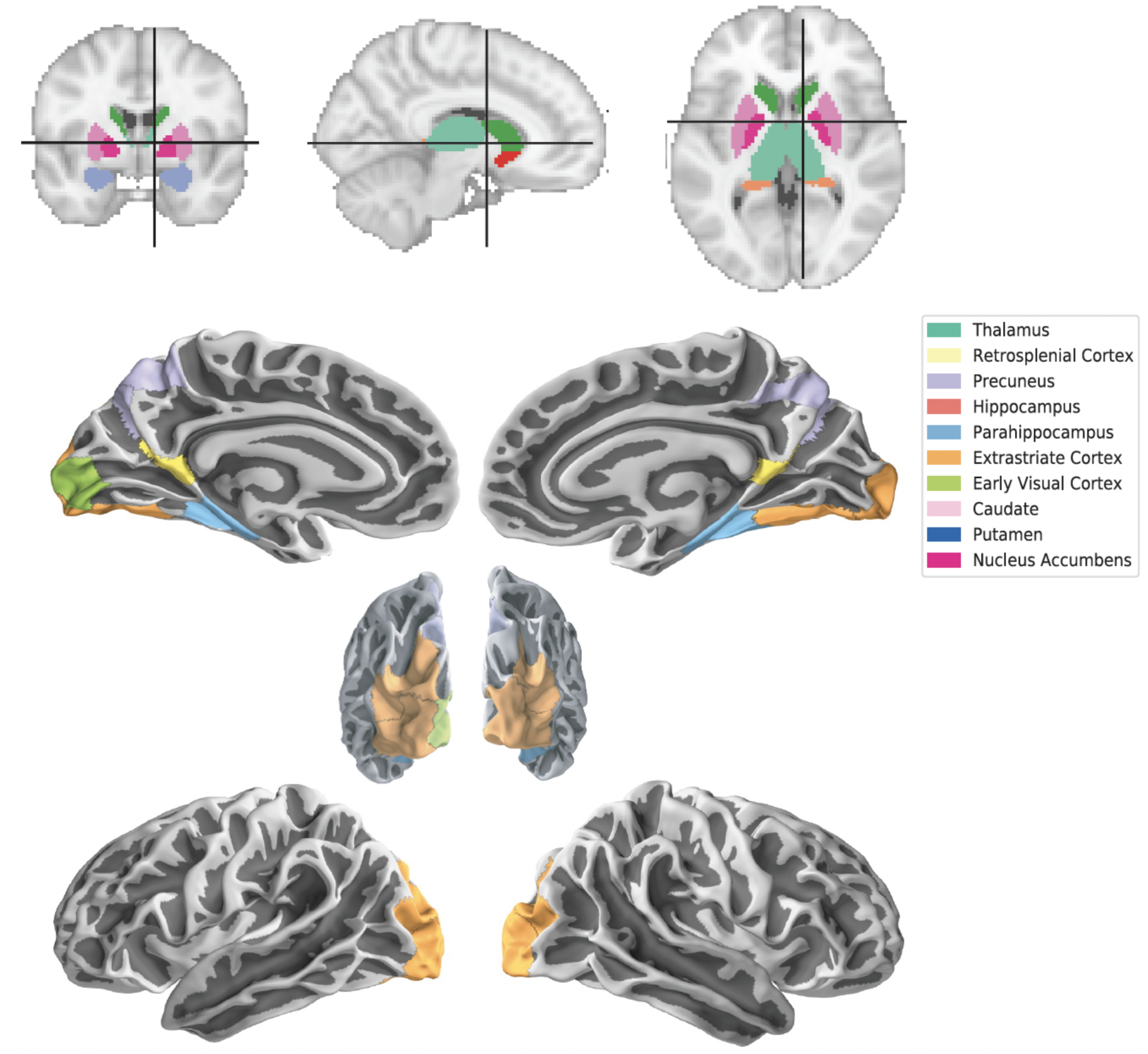
Regions of interest (ROIs) for the analysis: thalamus, retrosplenial cortex, precuneus, hippocampus, parahippocampal cortex, extrastriate cortex, early visual cortex, caudate, putamen, nucleus accumbens. All cortical ROIs (retrosplenial cortex, precuneus, extrastriate cortex, parahippocampal cortex, and early visual cortex) were generated for each participant’s brain in the 521MNI space based on the Schaefer 100 regions atlas. All subcortical ROIs (including thalamus, hippocampus, caudate, putamen, and nucleus accumbens) were generated for each participant’s brain in the 521MNI space based on the Harvard-Oxford subcortical probabilistic atlas with 25% threshold. All ROIs included bilateral regions.

**Figure S2.**
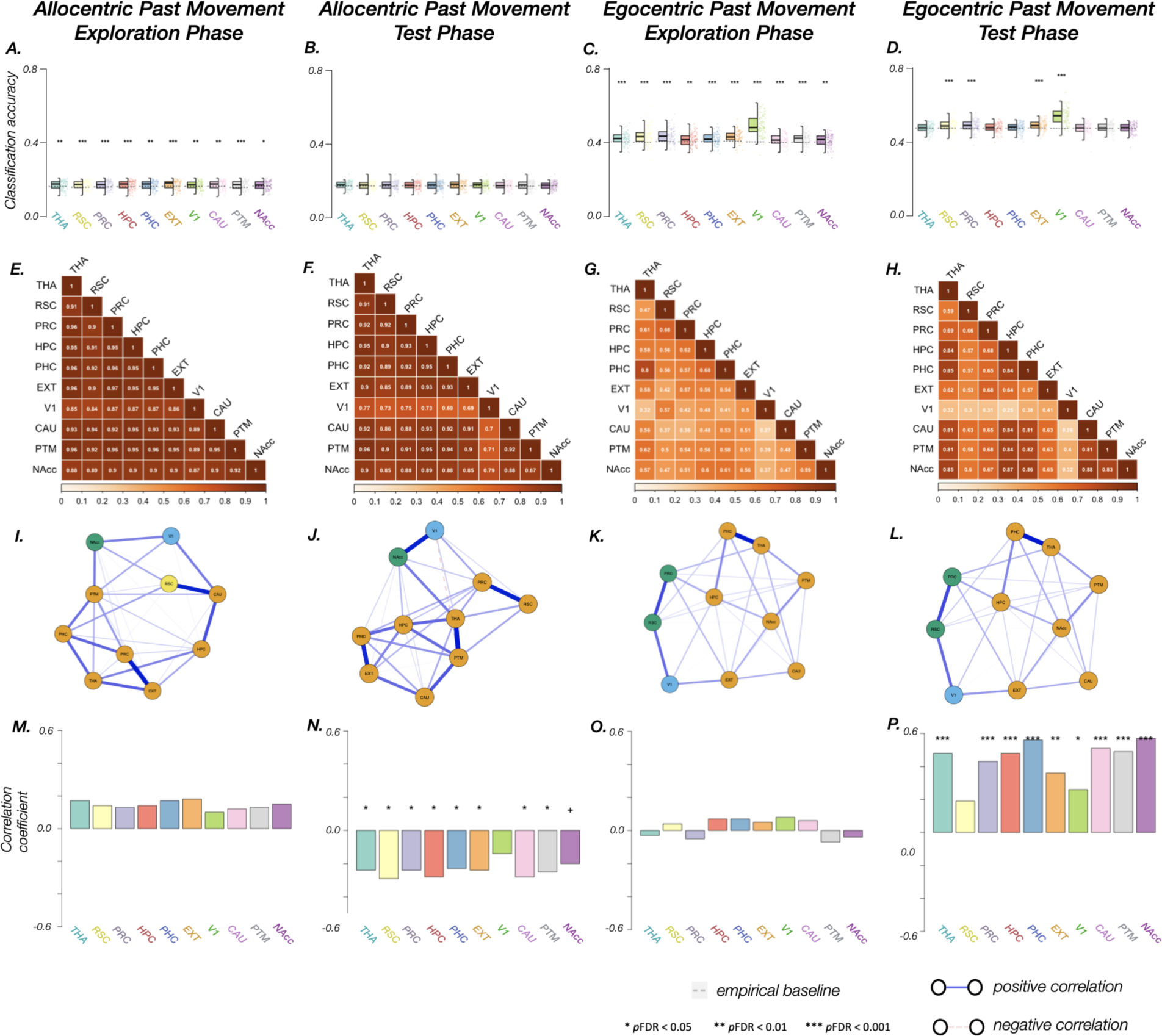
Performance of allocentric and egocentric past movement in 10 ROIs (THA, thalamus; RSC, retrosplenial cortex; PRC, precuneus; HPC, hippocampus; PHC, parahippocampal cortex; EXT, extrastriate cortex; V1, early visual cortex; CAU, caudate; PTM, putamen; NAcc, nucleus accumbens). These classifications are when the participant is stationary at an intersection, but we classified based on their previous allocentric or egocentric movement. A.-D. Classification accuracy in each ROI during the four sections of the task: A. allocentric past movement during the exploration phase, B. allocentric past movement during the test phase, C. egocentric past movement during the exploration phase, D. egocentric past movement during the test phase; E.-H. Pearson correlation matrices of classification accuracy in each pair of ROIs in the four sections of the task; I.-L. Correlation networks among ROIs in the four sections of the task. Each color represents a separate subnetwork. note: for better visualization, network connections were tuned with Gaussian Markov random field estimation using graphical LASSO and extended Bayesian information criterion to select optimal regularization parameters. As a result, weak Pearson correlations in correlation matrices may appear as negative connections in network visualization. Nodes represent ROIs and edges represent tuned correlation coefficients. Subnetworks were generated based on hierarchical clustering and the optimal number of clusters were determined with silhouette score; M.-P. Correlation coefficients between navigation performance (i.e., accuracy) and classification accuracy in each ROI in the four sections of the task.

**Figure S3.**
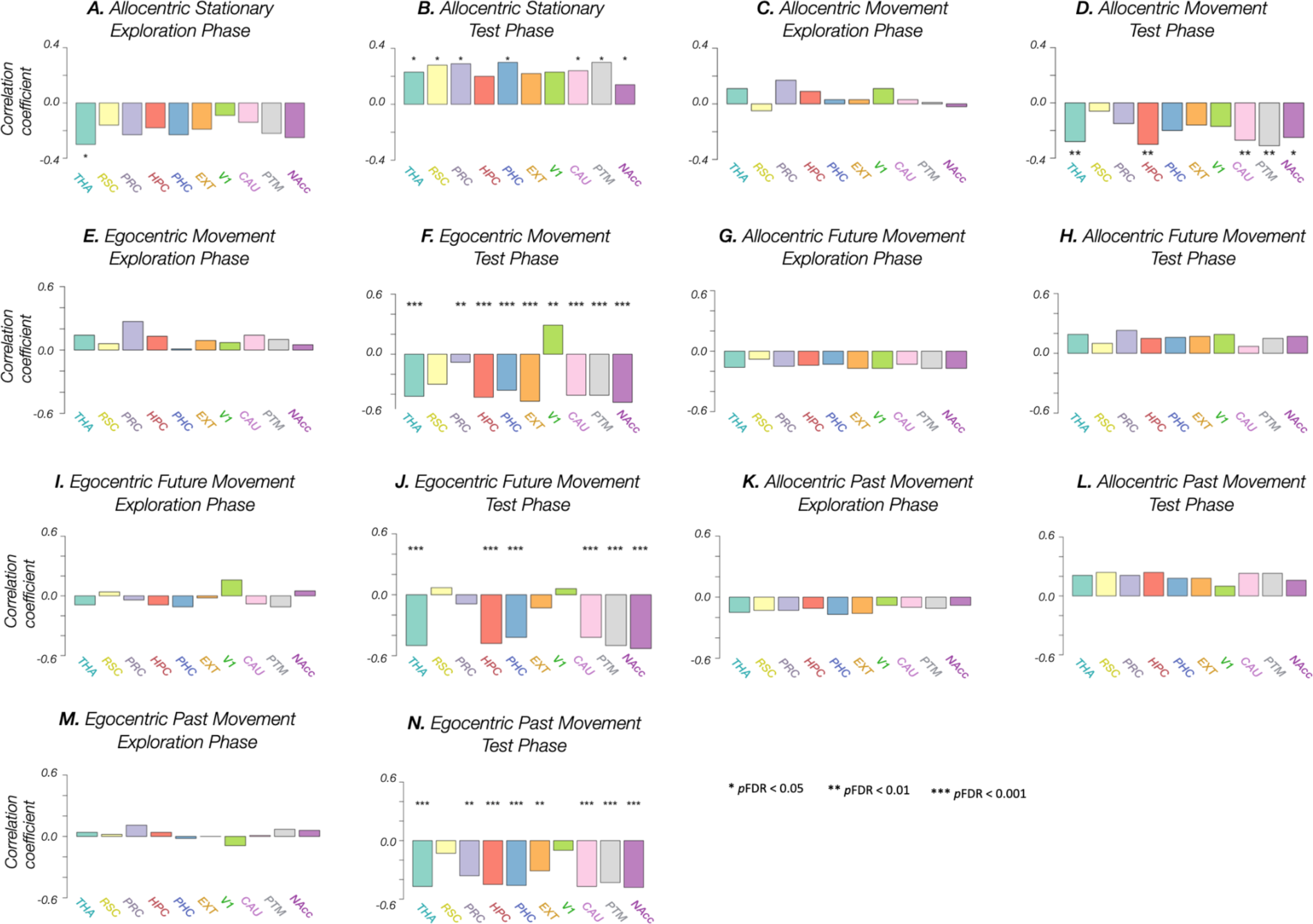
Correlation coefficients between excess path ratio (path efficiency) and classification accuracy in 10 ROIs (THA, thalamus; RSC, retrosplenial cortex; PRC, precuneus; HPC, hippocampus; PHC, parahippocampal cortex; EXT, extrastriate cortex; V1, early visual cortex; CAU, caudate; PTM, putamen; NAcc, nucleus accumbens). Excess path ratio is the length of the path traveled divided by the shortest ideal path, for correct trials only. Thus, a lower score is a more efficient path, so negative correlations indicate that people with more efficient paths had better classification accuracy. A. allocentric stationary during the exploration phase, B. allocentric stationary during the test phase, C. allocentric movement during the exploration phase, D. allocentric movement during the test phase, E. egocentric movement during the exploration phase, F. egocentric movement during the test phase, G. allocentric future movement during the exploration phase, H. allocentric future movement during the test phase, I. egocentric future movement during the exploration phase, J. egocentric future movement during the test phase, K. allocentric past movement during the exploration phase, L. allocentric past movement during the test phase, M. egocentric past movement during the exploration phase, N. egocentric past movement during the test phase

